# LAD: an R package to estimate leaf angle distribution from measured leaf inclination angles

**DOI:** 10.1101/2022.10.28.513998

**Authors:** Francesco Chianucci, Lorenzo Cesaretti

## Abstract

Leaf angle distribution (LAD) is an important factor for characterizing the optical features of vegetation canopies. The characterization of LAD requires direct measurements of leaf inclination angles, which can be obtained from manual clinometer measurements or leveled digital photography.

Package `LAD` allows to calculate the Leaf angle distribution (LAD) function and the G-function from measured leaf inclination angles. Using the package, the LAD distribution and G-function is derived by fitting a two parameter Beta distribution. Summary leaf angle statistics and distribution type is also calculated, by comparing the obtained LAD against theoretical distribution by de Wit (1965).

## Introduction

Leaf angle distribution (LAD) is an important factor for characterizing the optical features of vegetation canopies (Ross 1981). It influences several processes such as photosynthesis, evapotranspiration, spectral reflectance and absorptance (Vicari et al. 2019). LAD influence on radiation transmission is also described by the leaf projection function (also known G-function), which is the projection coefficient function of unit foliage area on a plane perpendicular to the viewing direction (Ross 1981). Both parameters are therefore crucial to indirectly estimate Leaf Area Index (LAI) from gap fraction or radiation trasmission measurements.

Despite its importance, LAD is one of the most poorly characterized parameters due to the difficulty of directly measure leaf inclination angles, particularly in tall trees. As alternative to direct manual measurements, Ryu et al. (2010) proposed a robust and simple leveled photographic method to measure leaf inclination angles, which was proven comparable to manual clinometer measurements (Pisek et al. (2011)). Using this leveled photographuic method, some dataset of leaf inclination angle distributions have been compiled by some authors (Pisek et al. 2013; Raabe et al. 2015; Chianucci et al. 2018; Pisek and Adamson 2020).

In this article we presented “LAD”, an R package to calculate the Leaf angle distribution (LAD) function and the G-function from measured leaf inclination angles.

## Basic theory

The method proposed by Ryu et al. (2010) consists of acquiring leveled images of the canopy using a digital camera. As some species may display phototropism, leaves shall be measured in all the azimuth directions and along the vertical profile of the surrounding canopy. Crowns of trees can be observed using towers, extendable poles, ladder, nearby tall buildings, or unmanned aerial vehicles (McNeil et al. 2016). The measurement of leaf inclination angle requires the identification of the leaf plane, from which the leaf normal is measured (Fig. 1). For this reason, the leaves must be selected from those oriented approximately parallel to the viewing direction of the camera (i.e., the leaves shown as a line in the image, Fig. 1), avoiding bended leaves to be measured.

**Figure 1.**
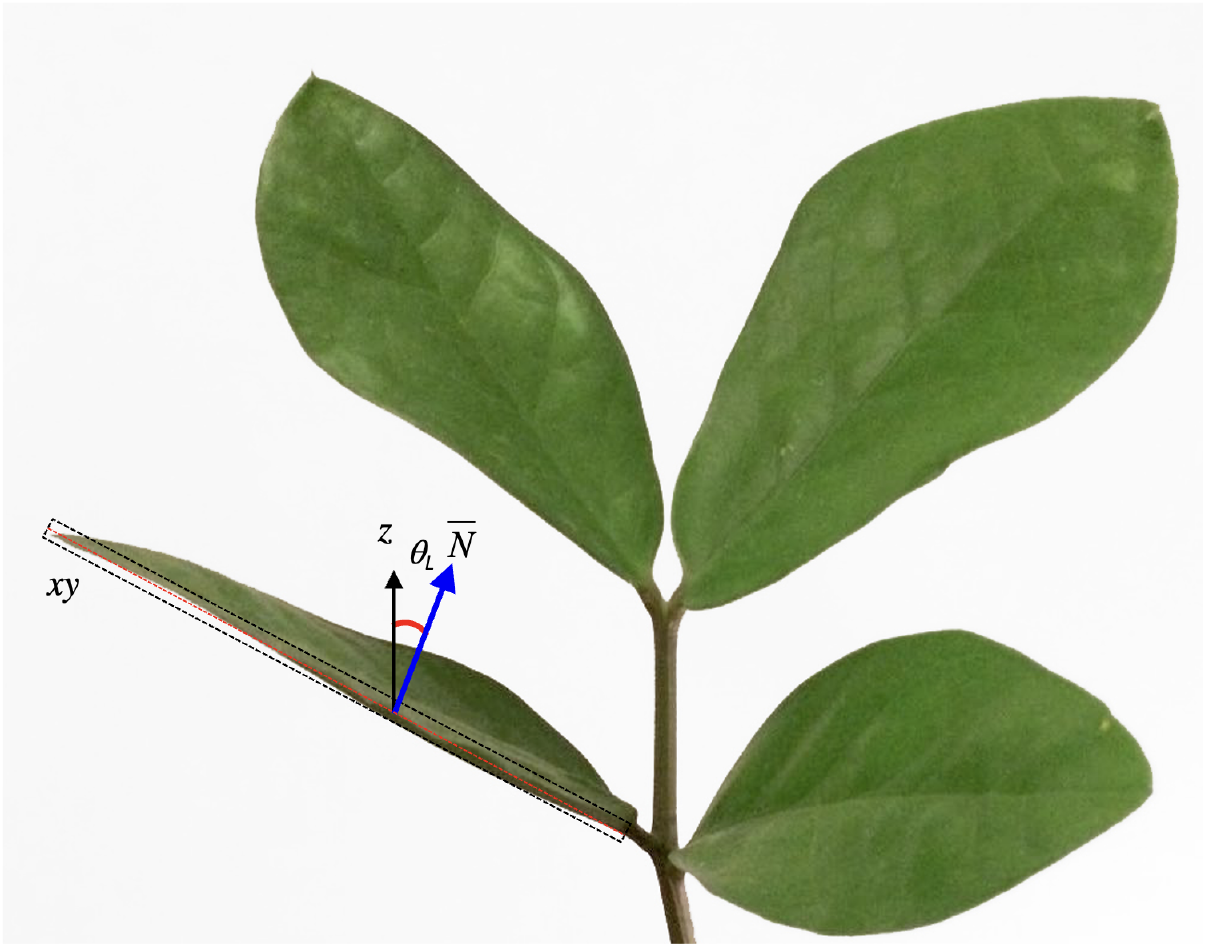
Example of leaf inclination angle measurements from leveled photography. The measure of leaf inclination angle requires the identification of a hypothetical leaf plane xy, from which the leaf surface normal 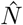 is calculated with respect to the zenith (z). In a 2-D space, such as a digital image, the only measurable leaf normal is that oriented perpendicular to the camera viewing direction, since the leaf inclination plane is parallel to the leaves (the red dashed line on the left side of the figure). From Chianucci et al. 2018, modified

Once a reliable set of leaf inclination angles measurements *θ*_*L*_ are taken (a minimum of 75 measurements per species are recommended by Pisek et al. (2013)), two parameters *µ, ν* are derived for fitting a Beta distribution:

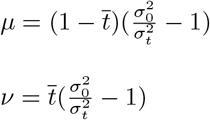

where 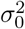 and 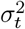 are the maximum standard deviation and variance of t, respectively, and are calculated following Wang et al.(2007):

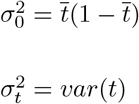

with *t* = 2*θ*_*L*_*/π*, and *θ*_*L*_ expressed as radians.

The two *µ, ν* parameters were then used to fit a Beta distribution function to represent the LAD of the considered canopy Goel and Strebel (1984):

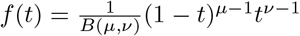

Following de Wit (1965), LAD functions can be described using six common functions based on empirical evidence of the natural variation of leaf normal distributions and mathematical considerations:

In spherical canopies, the relative frequency of leaf inclination angle is the same as for a sphere; planophile canopies are dominated by horizontally oriented leaves; plagiophile canopies are dominated by inclined leaves; erectophile canopies are dominated by vertically oriented leaves; extremophile canopies are characterized by both horizontally and vertically oriented leaves; uniform canopies are characterized by equal proportion of leaf inclination angles for any angle.

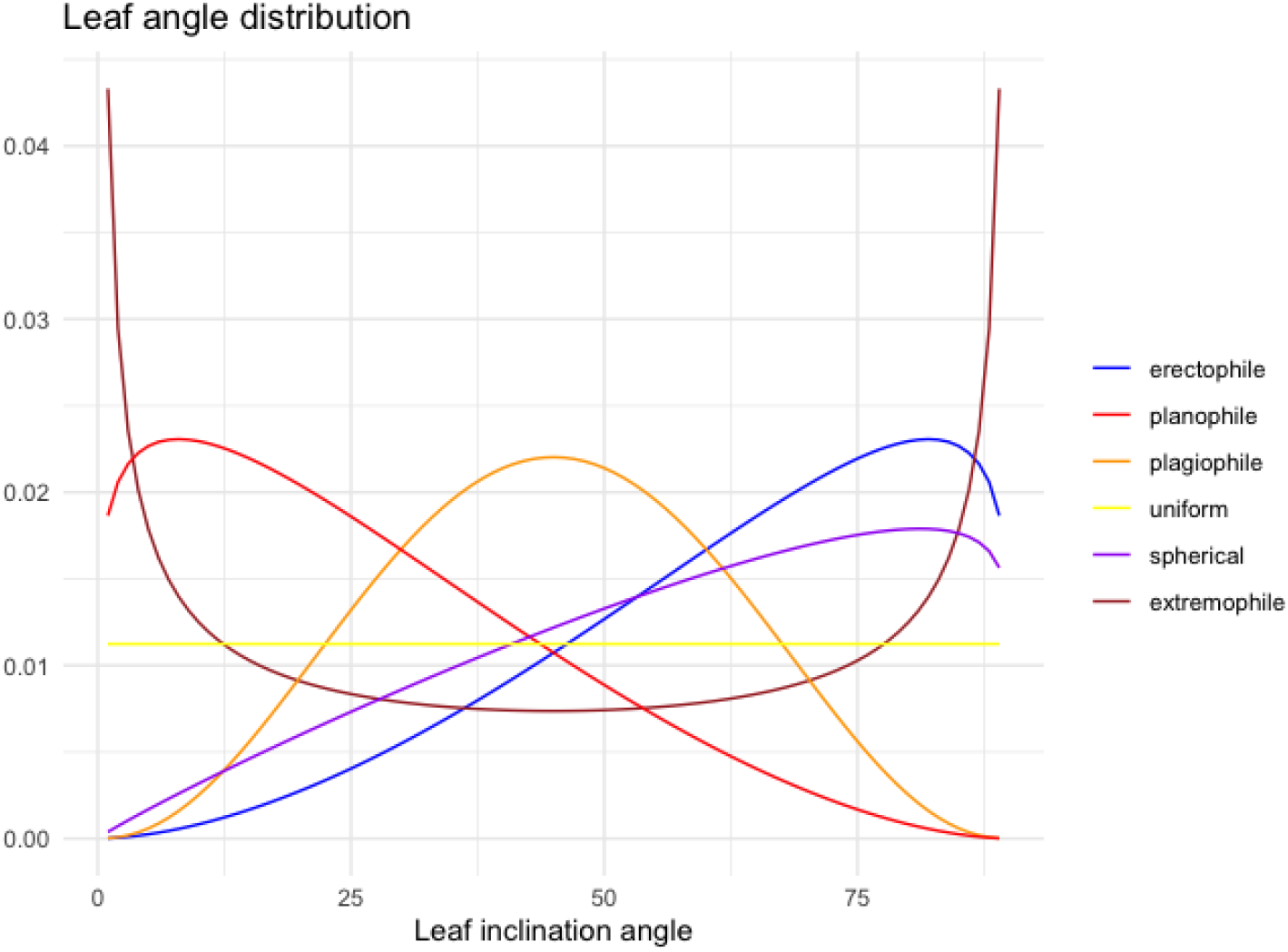

The fitted LAD function can then be compared with these six distributions, to determine the actual distribution type. The distribution type is classified using a leaf inclination index (Ross (1975)):

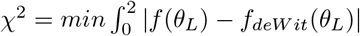

The LAD function can also be used to calculate the G-function (Ross (1981)), which corresponds to the fraction of foliage area on a plane perpendicular to the view angle *θ*:

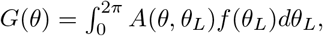

where:

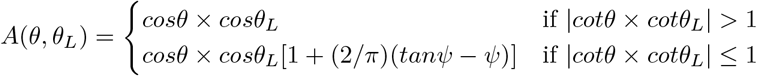

and:

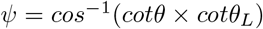

### Installation

You can install the development version of LAD from GitLab using `devtools` (Wickham et al. 2021):

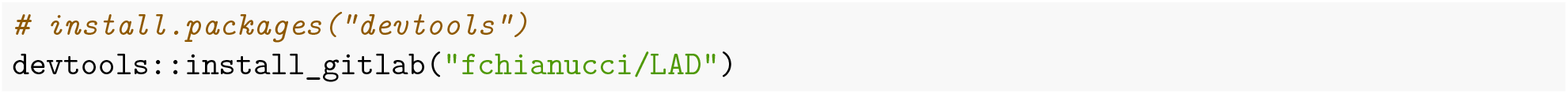

## The LAD package

The package have two key functions to calculate LAD, G-function, summary leaf inclination angle statistics:

### fitLAD(): fit a LAD distribution

fitLAD calculates the LAD function and G-function from two-parameters *µ, ν* of Beta distribution. The functions returns a list with two elements:

- the first element (*‘dataset’*) is a dataframe with three columns indicating the fitted LAD function (*pdf*), the G-function (*G*), for view or inclination angle (*theta*);
- the second element (*‘distribution’*) is a vector containing the matched distribution type;
- the extra argument `plot` allows to display the calculated LAD and G from measured leaf inclination angles.

Example of graphical output:

~~~
fitLAD(4,2,plot=T)
~~~

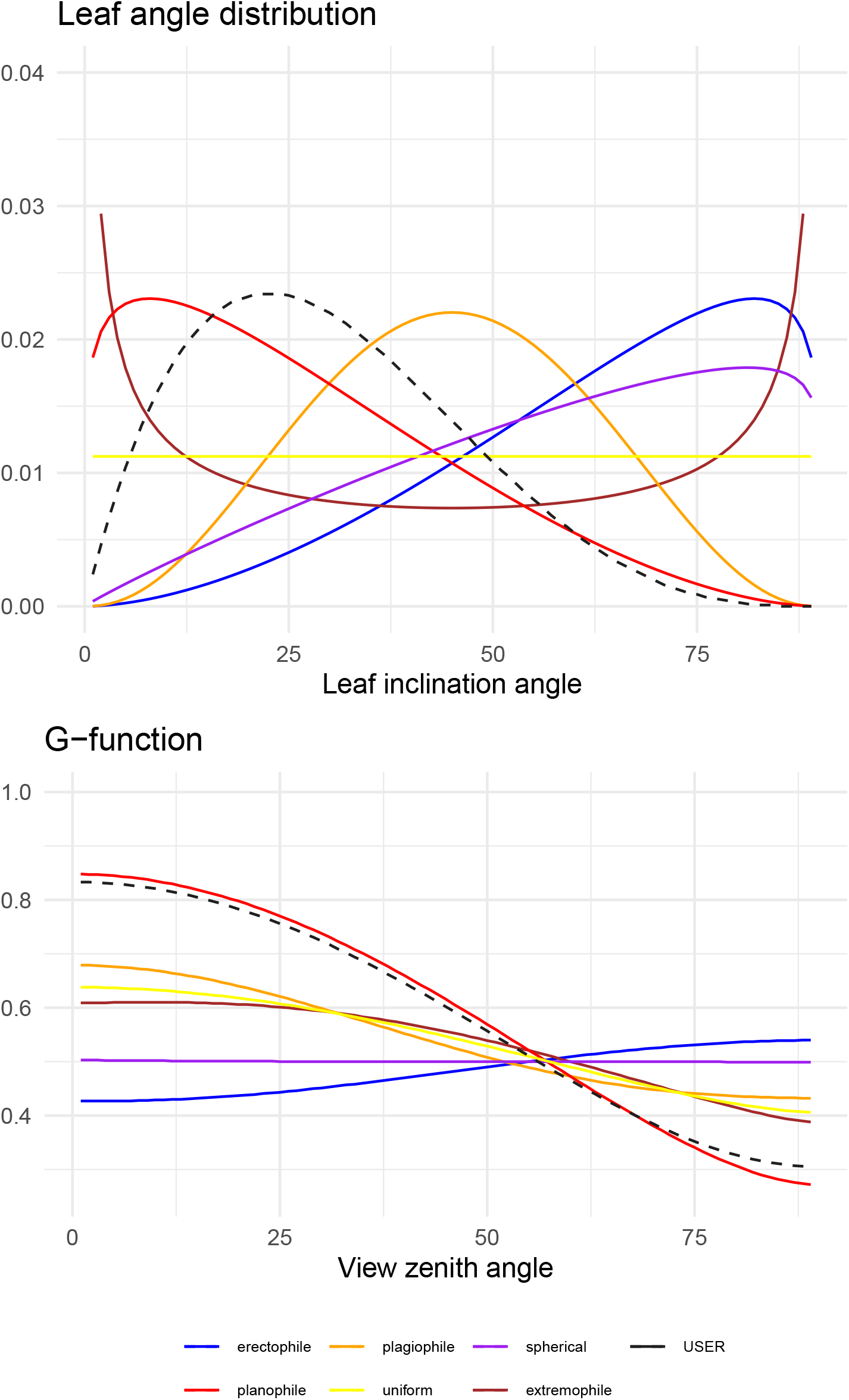

### calcLAD(): calculate LAD statistics

The `calcLAD(dataframe,angles,type,…)` function allows to calculate either summary leaf inclination angle statistics or the LAD function and G-function from leaf inclination angle measurements. Three important considerations must be drawn when using this function:

1. The function considers a *dataframe* as input; it doesn’t work on single vector(s) of measurements;
2. The function needs one or more *grouping* column(s) to calculate the attributes;
3. The function yields a *summarized* list of attributes derived from measured leaf inclination angles or an *extended* calculation of LAD and G-function depending of the function argument `type`.

### Package usage

We will illustrate the function usage with example data from Chianucci et al. (2018), which represents the largest leaf inclination angle dataset for 138 temperate and boreal woody species.

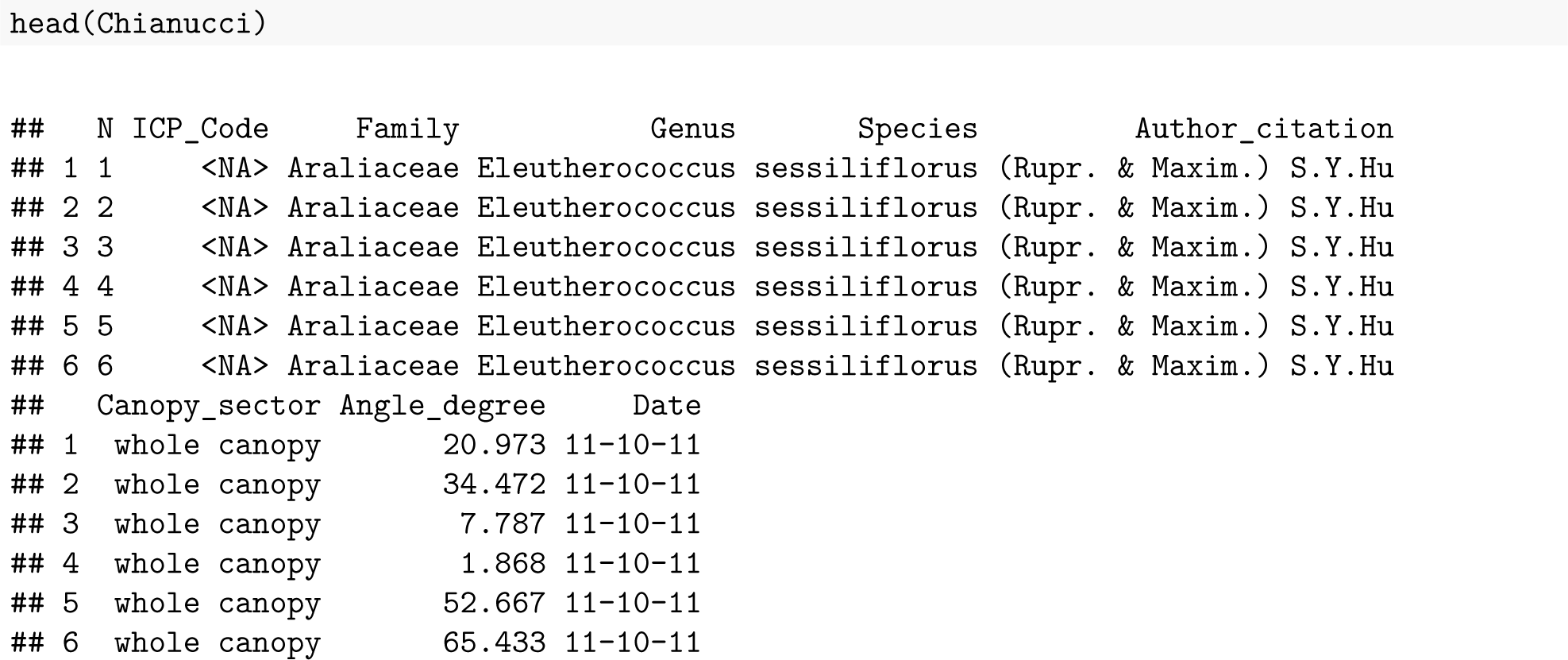

The `calcLAD()` function requires selecting the columns with the leaf inclination angle measurements data (in degrees), the type of calculations to be performed (*‘summary’* or *‘extended’*) and the grouping column(s).

By selecting `type=‘extended’` the function calculate LAD function (*pdf*) and G-function (*G*) for each group:

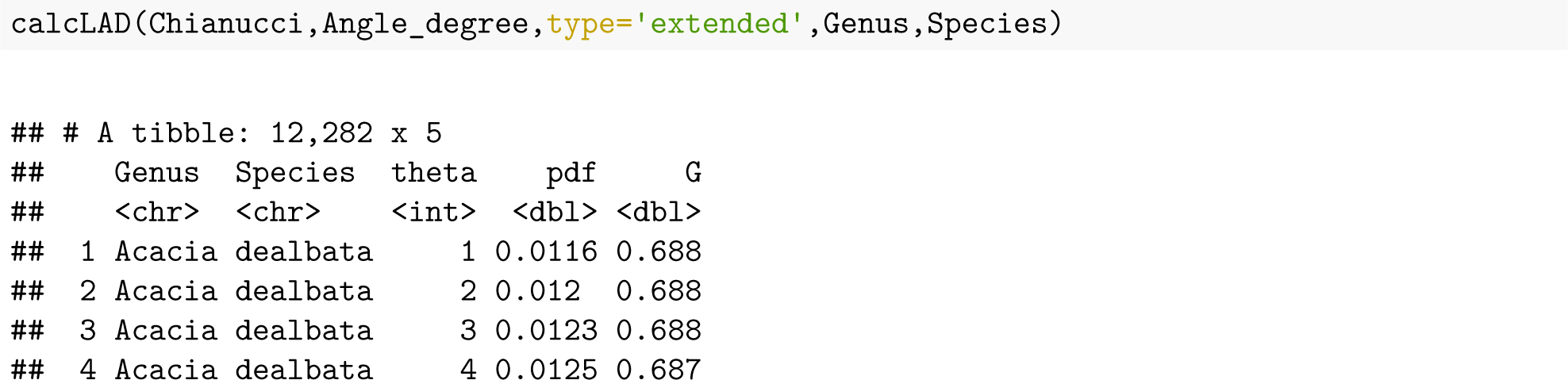
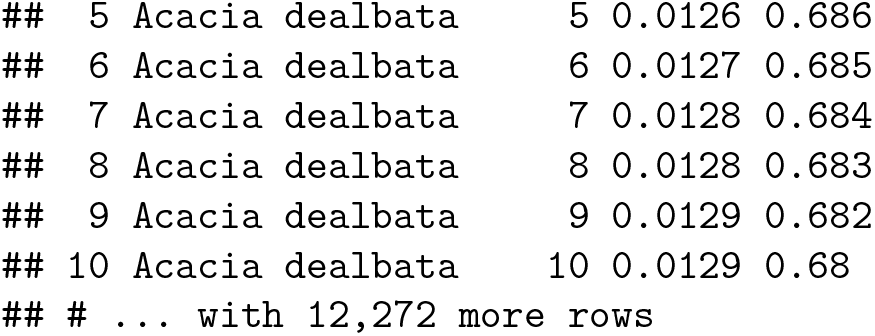

By selecting `type=‘summary’` the function calculate mean statistics (mean leaf inclination angle (MTA); standard deviation (SD), frequency (NR), Beta parameters (mu,nu) and distribution type):

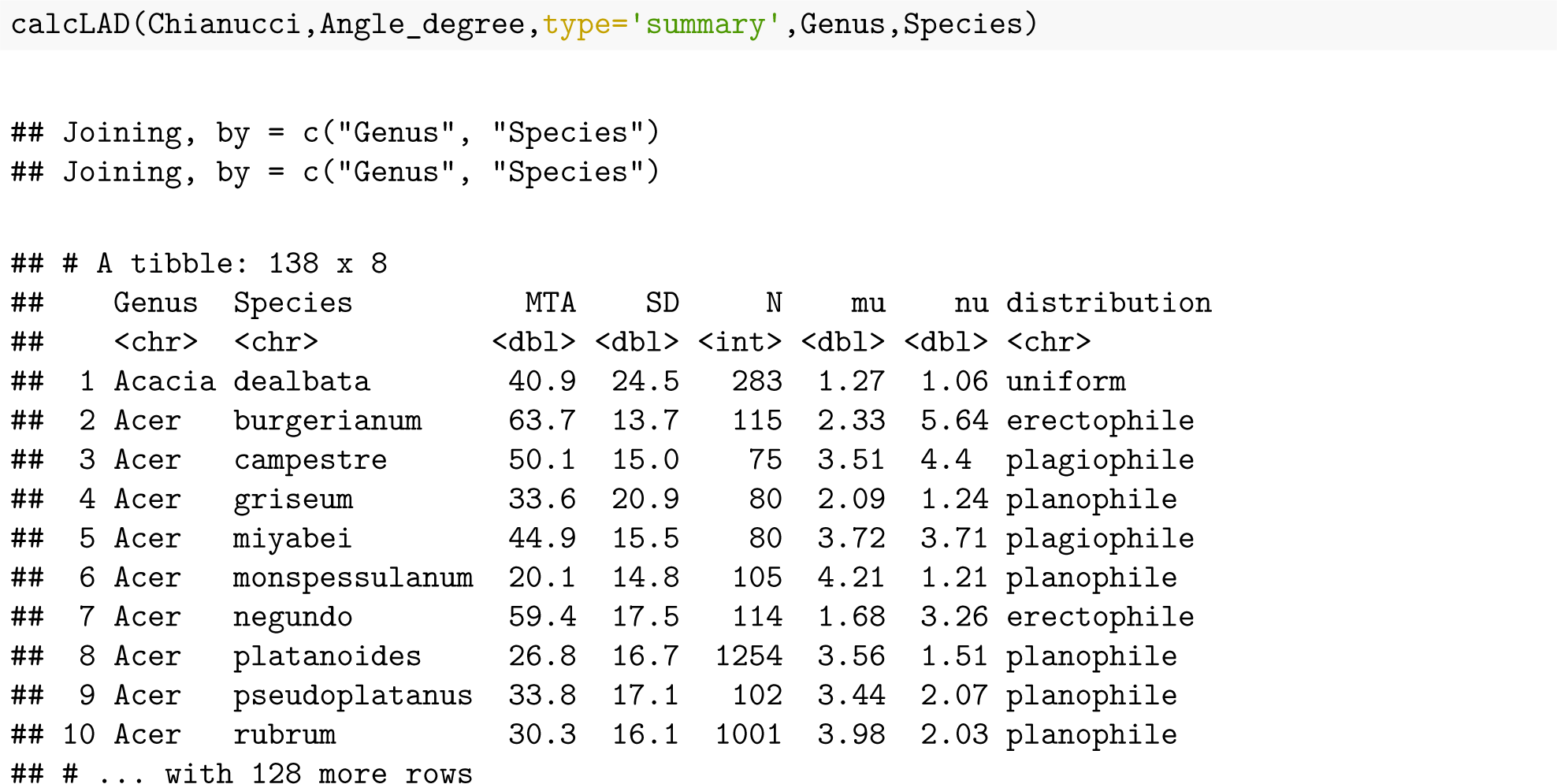

